# Unfolded Protein Response Factor ATF6 Augments T Helper Cell Responses and Promotes Mixed Granulocytic Airway Inflammation

**DOI:** 10.1101/2023.03.06.531397

**Authors:** Dandan Wu, Xing Zhang, Kourtney M. Zimmerly, Ruoning Wang, Chunqing Wang, Xiang Wu, Meilian Liu, Xuexian O. Yang

## Abstract

The unfolded protein response (UPR) is associated with the risk of asthma, including treatment-refractory severe asthma. Several recent studies demonstrated a pathogenic role of activating transcription factor 6α (ATF6α or ATF6), one of the essential arms of UPR, in airway structural cells. However, its role in T helper (TH) cells has not been well examined. In this study, we found that ATF6 was selectively induced by STAT6 and STAT3 in TH2 and TH17 cells, respectively. ATF6 upregulated UPR genes and promoted the differentiation and cytokine secretion of TH2 and TH17 cells. T cell-specific *Atf6*-deficiency impaired TH2 and TH17 responses *in vitro* and *in vivo* and attenuated mixed granulocytic experimental asthma. ATF6 inhibitor Ceapin A7 suppressed ATF6 downstream gene expression and TH cell cytokine expression in both murine and human memory CD4^+^ T cells. At the chronic stage of asthma, administration of Ceapin A7 lessened TH2 and TH17 responses *in vivo*, leading to alleviation of both airway neutrophilia and eosinophilia. Thus, our results demonstrate a critical role of ATF6 in TH2 and TH17 cell-driven mixed granulocytic airway disease, suggesting a novel option to combat steroid-resistant mixed and even T2-low endotypes of asthma by targeting ATF6.

## Introduction

Asthma is a chronic lung disease that inflames and constricts the airways, leading to breathing difficulty. Among all cases of asthma, about a half of cases are CD4^+^ T helper type 2 cell (TH2 or T2)-biased ^1^, manifesting an atopic phenotype including production of IgE to environmental antigens, airway eosinophilia, and expression of TH2 cytokines such as IL-4, IL-5, and IL-13. In contrast to the eosinophilic T2 high endotype, the non-eosinophilic T2-low endotype is more dominant in patients with severe asthma ^2^, and displays a neutrophilic phenotype with prominent expression of TH17 type cytokines such as IL-17 (also termed IL-17A) and IL-17F ^3–6^. T2 low asthma is treatment-refractory and associated with a high risk of mortality and disability. Airway colonization of bacteria (such as *Haemophilus influenza*, *Tropheryma whipplei*, *Streptococcus pneumonia*, and *Moraxella catarrhalis*) ^7, 8^ and fungi (such as *Alternaria*, *Aspergillus*, *Cladosporium,* and *Penicillium*) ^9^ induces TH17 responses and leads to airway neutrophilia ^9, 10^. Besides T2 high and T2 low endotypes, the eosinophilic and neutrophilic mixed granulocytic endotype contains both TH2 and TH17 signatures ^11^. *C. albicans* airway infection elicits protective TH (including TH17 and TH2) responses in murine ^12^, whereas uncontrolled fungal colonization in the airway may lead to mixed or even TH17-biased asthma. In summary, the distribution of granulocytes among different endotypes parallels the development of TH2 and TH17 cells. However, the underlying mechanism has not been well understood.

TH cells (and also immune infiltrates) belong to secretory cells. When receiving stimuli from T cell receptor (TCR) engagement, costimulation, and cytokine signals, TH cells produce massive amounts of effector molecules, including their signature cytokines, such as TH2 cytokines (IL-4, IL-5, and IL-13) and TH17 cytokines (IL-17A, IL-17F, IL-22, and GM-CSF). During this process, the accumulation of misfolded/unfolded proteins in the endoplasmic reticulum (ER) leads to ER stress; ER stress further activates UPR pathways: activating transcription factor 6α (ATF6α, or ATF6 hereafter), inositol-requiring enzyme 1 (IRE1), and PKR-like ER kinase (PERK). Among these, ATF6 and IRE1, after activation, regulate X-box protein 1 (*Xbp1*) gene transcription and splicing (to functional *Xbp1s* mRNA), respectively ^13–15^. The transcription factor XBP1s induces a set of ER-associated proteins, including BiP (HSPA5 or GRP78), ATF6, ERdj4, p58IPK, EDEM, RAMP-4, PDI-P5, and HEDJ ^16^, and is essential in the secretory function of cells, including immune cells ^13, 14, 17^. XBP1s also promotes autophagy through induction of Beclin-1 ^18^. In addition to the *Xbp1* gene, activated ATF6 (the N-terminal fragment, or ATF6f) transactivates the genes encoding P58IPK, ERdj4, BiP, GRP94, and CHOP ^16, 19^.

Besides, ATF6f suppresses UPR-induced apoptotic program and facilitates the recovery from acute stress ^20^. Taken together, the IRE1 and ATF6 pathways of UPR expand the capacity of ER in protein folding, maturation, and secretion, resolve ER stress and restore cellular homeostasis.

The UPR driven by protein synthesis-induced ER stress is involved in many conditions including lung diseases ^21, 22^. For instance, IRE1β, the mucosal epithelial cell-specific form of IRE1, is required for goblet cells to produce mucins; however, in experimental asthma, IRE1β promotes goblet cell metaplasia ^21^. Airway epithelial cells from asthmatic patients display an increased level of ER stress factor ERp57 ^23^. In *Aspergillus fumigatus*-induced murine asthma, the lung tissue expresses increased levels of UPR markers, including BiP, CHOP, p-IRE1α, p-eIF2α, XBP1, and ATF4 ^24^. Similarly, house dust mite (HDM) provokes a vigorous ER stress-UPR reaction in human and murine airway epithelial cells, including the induction of ATF6 ^25^. GWAS studies have associated ER stress factor orosomucoid-like (ORMDL) 3 with the risk of asthma^26–28^, including severe asthma ^29, 30^. ORMDL3 directly interacts with sarco-endoplasmic reticulum calcium ATPase 2b (SERCA2b) and inhibits calcium (Ca^2+^) influx into the ER, leading to UPR activation ^28^. ORMDL3 promotes airway smooth muscle hyperplasia *in vitro* ^31^. In an *Alternaria* asthma model, ORMDL3 promotes T2-high airway disease through activation of the ATF6 arm of UPR in airway epithelial cells ^32^. Human airway epithelial brushings from asthmatic subjects also demonstrate upregulation of several ATF6-related transcripts compared with those from healthy controls ^33^. It is not clear whether the ORMDL3-ATF6 axis also regulates UPR in TH cells and contributes to the development of airway inflammation as that of epithelial cells.

In the current study, we found that ATF6 is selectively expressed by TH2 and TH17 cells in a signal transducer and activator of transcription 6 (STAT6) and STAT3-dependent manner. *In vitro*, *Atf6*-deficiency decreases the expression of ER stress factors and impairs the differentiation and cytokine production of TH cells. *In vivo*, both genetic ablation and pharmaceutical inhibition of ATF6 lead to alleviated TH2 and TH17 responses and mixed granulocytic airway inflammation. These data demonstrate that ATF6 mediates ER stress and UPR in TH cells, promotes airway inflammation, and subsequently, aggravates both eosinophilic and neutrophilic airway inflammation.

## Results

### Upregulation of ATF6 in T cells and asthmatic lungs

Heightened UPR is associated with human asthma. The ORMDL3-ATF6 pathway drives UPR in asthmatic airway epithelial cells and promotes experimental T2-high airway disease ^32^.

Consistently, in a house dust mite-*Aspergillus fumigatus* (HDM-AF)-induced mixed type asthma model, we observed that the mRNAs encoding ATF6 and its downstream genes *Xbp1u* (unspliced form of *Xbp1*), *Xbp1s*, *Hspa5* (encoding BiP), and *Ddit3* (encoding CHOP) were highly induced in the asthmatic lungs compared with those from non-immunized mice (Fig. 1A). Since a balance of TH2 versus TH17 responses governs an eosinophilic or neutrophilic or mixed phenotype, we asked whether ATF6-mediated UPR also plays a role in TH cells during asthmatic reactions. Upon activation and differentiation, TH cells commit to different subsets with specific effector functions. We found that during naïve CD4^+^ T cell activation with plate-bound anti-CD3 and anti-CD28, the transcription of *Atf6* gene was upregulated as early as 2 h and persistent up to 24 h (Fig. 1B). In a gene expression profiling analysis of *in vitro*- differentiated TH1, TH2, TH17, and regulatory (Treg) cells, we found that the expression of *Atf6* mRNA was elevated in TH1, TH2, TH17, and Treg cells compared to naïve CD4^+^ T cells, but the increases were more profound in TH2 and TH17 cells (Fig. 1C), suggesting a role of ATF6 in TH2 and TH17 cell function. We then generated T-specific *Atf6*-deficient (*Atf6*^fl/fl;Cd4Cre^; termed *Atf6* KO) mice by crossing *Atf6*^fl/fl^ mice with Cd4Cre ^34, 35^, in which ATF6 protein and mRNA could not be detected in TH cells (Fig. 1D). The memory CD4^+^ T cells from *Atf6* KO mice had decreased expression of XBP1s compared with those from *Atf6*-sufficient littermate control mice [*Atf6*^fl/fl^; termed *Atf6* wild-type (WT)] (Fig. 1D). Concordantly, *Atf6* KO memory CD4^+^ T cells expressed decreased mRNA levels of *Xbp1u*, *Xbp1*s, *Hspa5*, and *Ddit3* (Fig. 1E). Interestingly, *Atf6*-deficiency led to decreased expression of TH2 cytokines IL-4, IL-5, and IL-13, and TH17 cytokine IL-17 by memory CD4^+^ T cells with a trend toward decreased levels of TH1 cytokine IFNγ (Fig. 1F). Taken together, ATF6 is selectively expressed by TH2 and TH17 cells, mediates UPR, and enhances the expression of cytokines in memory TH cells.

**Figure 1.**
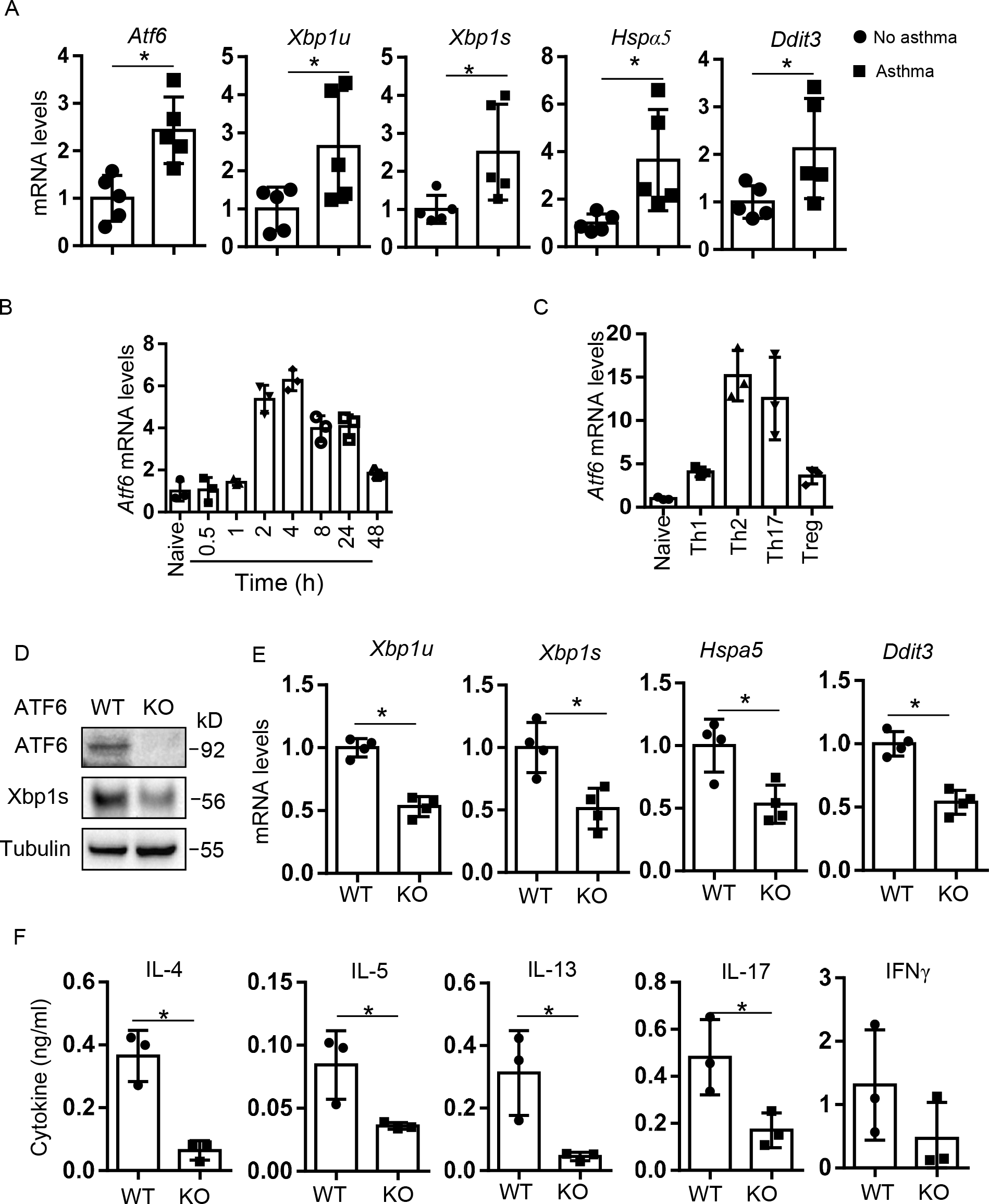
Expression of ATF6 in TH cells and in asthmatic lungs. (A) RT-qPCR of mRNA expression of *Atf6*, *Xbp1u*, *Hspα5*, and *Ddit3* in lungs from asthmatic and healthy mice. (B) *Atf6* mRNA expression by naive CD4^+^ T cells during activation. (C) *Atf6* mRNA expression by TH1, TH2, TH17, and inducible Treg cells. (D-F) Characterization of memory CD4^+^ T cells from *Atf6* WT and KO mice after activation with plate-bound anti-CD3 and anti-CD28. (D) Western Blot of ATF6 and XBP1s. Tubulin was used as a loading control. (E) mRNA expression of *Atf6*, *Xbp1u*, *Hspα5*, and *Ddit3*. (F) ELISA of TH cytokines IL-4, IL-5, IL- 13, IL-17, and IFNγ. *Actb* was used as a loading control (A-C, E). Data (mean ± SD) shown are a representative of 2 experiments (A, D) or a combination of 3 or 4 experiments (B, C, E, F). Student’s *t*-test, \**p* < 0.05.

### STAT proteins induce ATF6 in TH2 and TH17 cells

ATF6 is selectively induced in TH2 and TH17 cells (Fig. 1C). However, it is unclear how ATF6 is regulated in those TH cells. STAT proteins are broadly involved in cytokines-mediated TH cell responses ^36, 37^. IL-4-STAT6 signaling is essential for TH2 cell differentiation, whereas IL-6, IL-21, and IL-23 activate STAT3 and govern TH17 cell polarization and maintenance. Although other STATs also play a role in TH2 and/or TH17 cells, we focused on STAT6 and STAT3, the major inducers of TH2 and TH17 cells, respectively. To understand how the STATs control ATF6 expression, we searched their binding sites in the promoter and the conserved non-coding sequences of the *Atf6* gene by comparing mouse and human genome sequences using the ECR Browser and its associated rVista program ^38, 39^. We identified 2 clusters of putative STAT3 and STAT6 binding sites in the promoter region (Fig. 2A). Over-expression of a hyperactive form of STAT6 (STAT6VT) resulted in increased *Atf6* mRNA expression in TH2 cells compared with that of a control vector (Fig. 2B). Similarly, forced expression of persistently active STAT3 (STAT3C) increased the expression of *Atf6* mRNA in TH17 cells relative to that of a control vector (Fig. 2C). Furthermore, we found that in chromatin immunoprecipitation (ChIP) assays, p- STAT6 bound to its putative single sites 1 and 4, dual sites 6+7 (sites 6 and 7 are located closely and were detected by the same pair of primers) in the *Atf6* promoter region in TH2 cells (Fig. 2D). In Th17 cells, p-STAT3 bound to its putative dual sites 1+2, 3+4, and single sites 5 and 8 and marginally to the other sites (Fig. 2E). These data suggest that STAT6 and STAT3 proteins, after activation, bind to the *Atf6* promoter and transactivate the *Atf6* gene in TH2 and TH17 cells, respectively.

**Figure 2.**
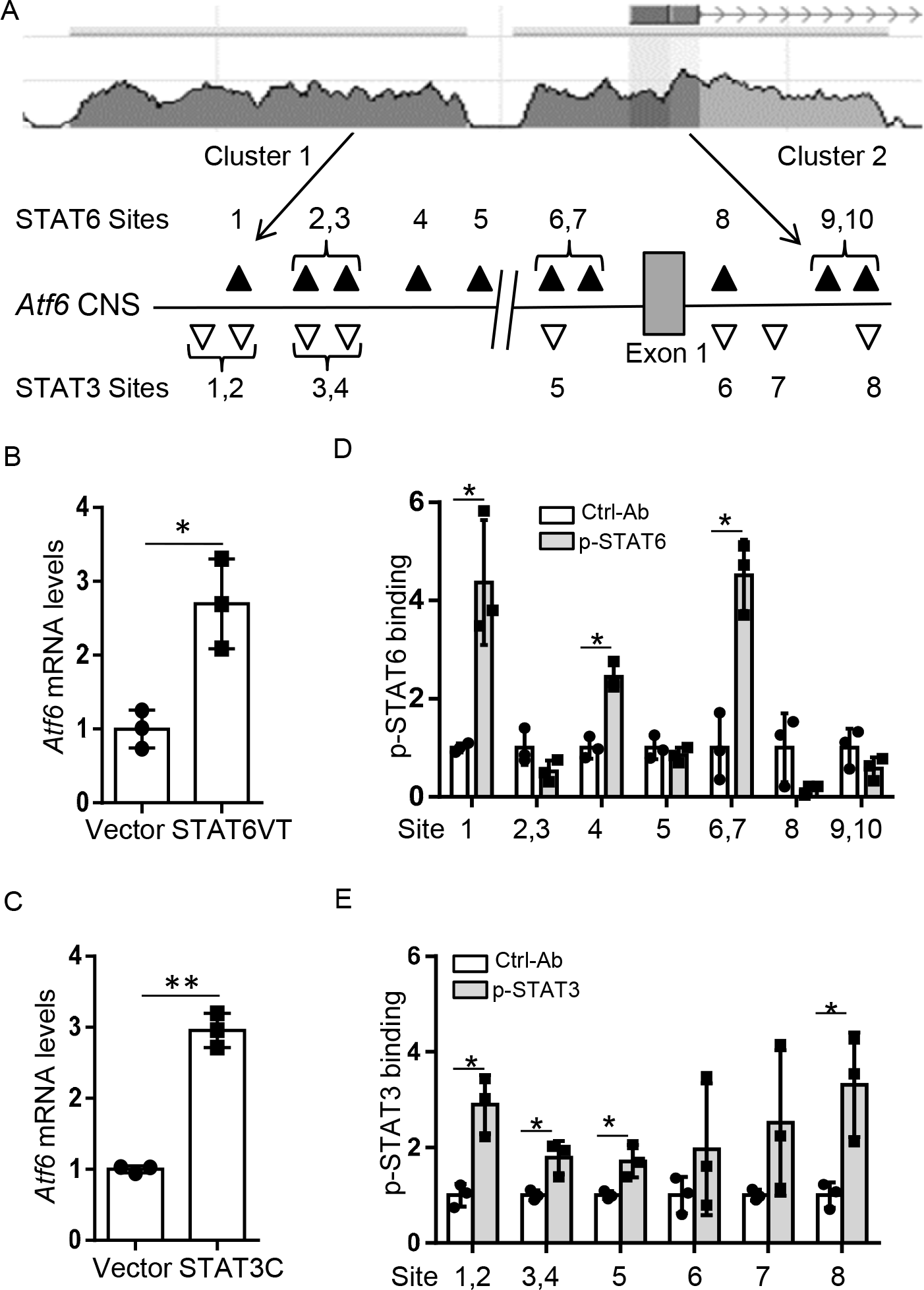
STAT3 and STAT6 induce ATF6 expression by TH2 and TH17 cells. (A) STAT3 and STAT6 binding sites conserved between humans and mice in the *Atf6* locus. (B) RT-qPCR of *Atf6* mRNA expression by TH2 cells with overexpression of STAT6VT or an empty vector. (C) *Atf6* mRNA expression by TH17 cells with forced expression of STAT3C or a control vector. (D) ChIP assays of pSTAT6 binding onto the *Atf6* locus in TH2 cells. (D) ChIP assays of pSTAT3 binding onto the Atf6 locus in TH17 cells. *Actb* was used as a loading control (B-C). Data are pooled from 3 experiments. Values are means and S.D. Student’s *t*-test, * *p* < 0.05.

### *Atf6*-deficiency impairs TH2 and TH17 responses *in vitro*

Since ATF6 is expressed during the early activation of naïve CD4^+^ T cells (Fig. 1B), we asked whether ATF6 plays a role in the initial lineage commitment of TH cells. To explore this, we skewed naïve CD4^+^ T cells from *Atf6* WT or KO mice under the TH1, TH2, TH17, and inducible Treg conditions, and assessed the resulting cells by using intracellular stain. As expected, *Atf6*- deficiency resulted in lower percentages of IFNγ^+^ TH1, IL-4^+^ TH2, and IL-17^+^ TH17 cells in the corresponding cultures (Fig. 3A-C). Under the inducible Treg condition, *Atf6* KO cells had only a marginal decrease in Foxp3 cell frequencies compared with *Atf6* WT cells (Fig. 3D). In addition to its role in TH cell differentiation, *Atf6*-deficiency impairs the expression of effector cytokines, IL-4, IL-5, and IL-13 by TH2 cells, IL-17 by TH17 cells, and in less extent, IFNγ by TH1 cells (Fig. 3E). Therefore, ATF6 promotes TH2 and TH17 polarization and cytokine production, implicating its role in TH2 and TH17-mediated mixed eosinophilic and neutrophilic disease.

**Figure 3.**
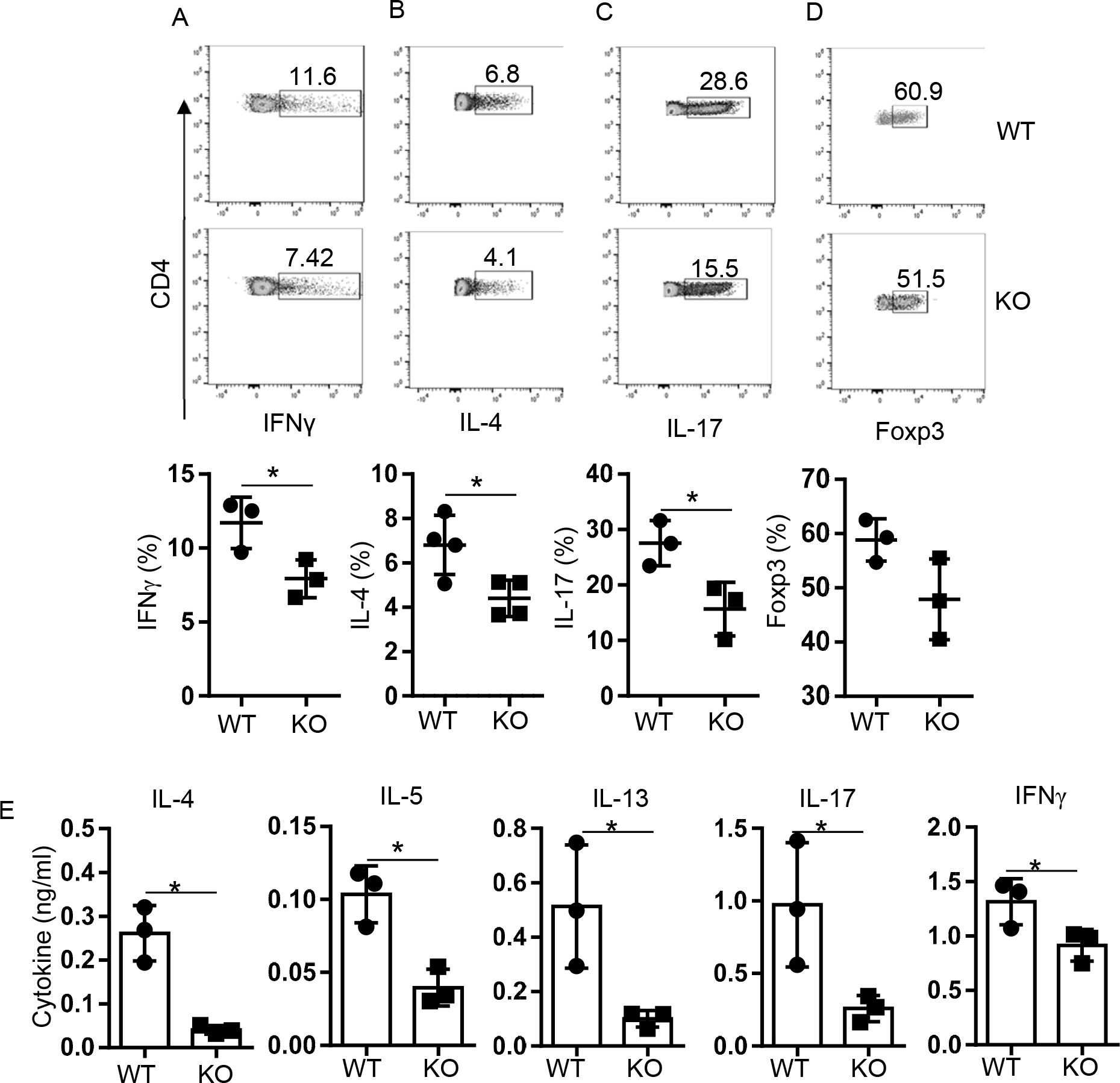
Atf6-deficiency impairs TH2 and TH17 differentiation and cytokine production. (A-D) Flow cytometry of *in vitro* polarized TH1, TH2, TH17, and inducible Treg cells following intracellular staining. (E) ELISA of cytokine expression by TH1, TH2, and TH17 cells.

### ATF6 mediates UPR in TH2 and TH17 cells

ATF6 regulates UPR factors and cytokine secretion in memory T cells (Fig. 1E-F). We next explored whether it also controls UPR in de novo differentiated TH cells and found that deletion of *Atf6* decreased the mRNA expression of ER stress factors *Xbp1u*, *Xbp1s*, *Hspa5*, and *Ddit3* in both TH2 and TH17 cells recently generated *in vitro* (Fig. 4A-B). In addition, *Atf6* KO TH2 and TH17 cells exhibited lower intensities of calreticulin (Calr), an ER-resident chaperon, which occupied less area in images acquired by imaging flow cytometry, compared with *Atf6* WT counterparts (Fig. 4C-D), suggesting that ATF6 is required for the expansion of ER. ATF6- mediated UPR is considered a post-transcriptional control. However, as a consequence of this post-transcriptional regulation, we observed that *Atf6*-deficiency impaired not only the secretion of TH2 and TH17 cytokines (Fig. 3E) but also the mRNA expression of those cytokines (Fig. 4E, G). Interestingly, we also observed a decrease of mRNAs encoding the master transcription factors GATA3 and RORγt [encoded by *Rorc(gt)*] for TH2 and TH17 cells, respectively (Fig. 4F, H). There is no evidence that ATF6 directly transactivates many TH2 and TH17 lineage-specific genes. To understand how ATF6 regulates the transcription in those TH cells, we examined the protein levels of GATA3 and RORγt in TH2 and TH17 cells, respectively, and found that *Atf6*- deficiency led to decreased levels of GATA3 and RORγt proteins in TH2 and TH17 cells, respectively (Fig. 4I-J). The result explains that the lessened protein levels of the master transcription factors cause impaired transcriptional activities in *Atf6* KO TH2 and TH17 cells compared with their WT counterparts. Therefore, ATF6 augments the ER capacity via induction of UPR factors and expansion of ER volume, and subsequently strengthens the transcription programs through upregulation of the master transcription factors, which together intensify the effector function of TH2 and TH17 cells (see the schematic in Fig. 4K).

**Figure 4.**
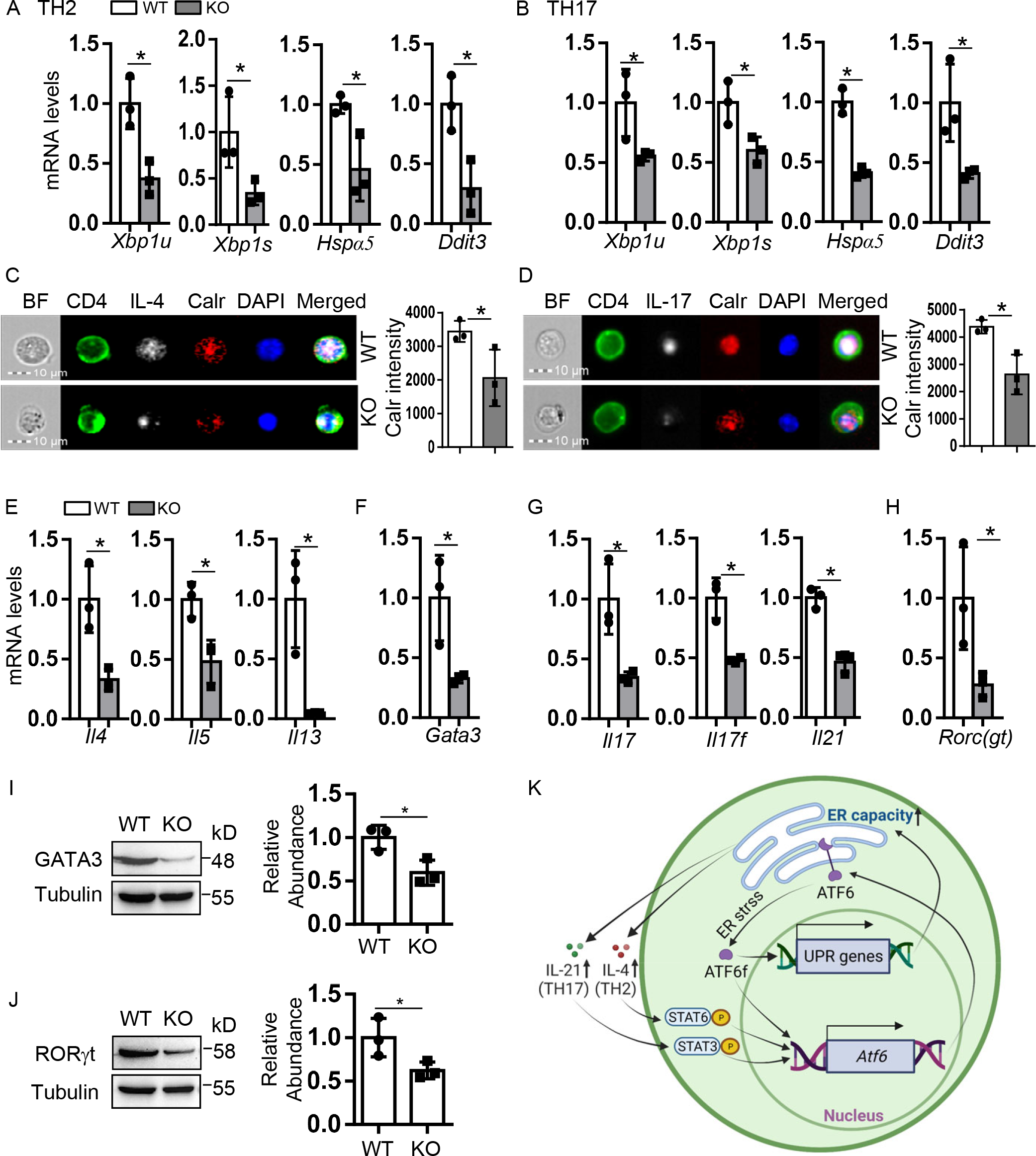
ATF6 regulates UPR and promotes TH2 and TH17 responses. (A-B) RT-qPCR of mRNA expression of ATF6 downstream genes in TH2 (A) and TH17 (B) cells with or without *Atf6*-deficiency. (C-D) Imaging flow cytometry of Calr in *Atf6* WT or KO TH2 (C) and TH17 (D) cells. Data points of Calr intensity were mean of 1000 CD4^+^ IL-4^+^ (C, right) or CD4^+^ IL-17^+^ (D, right) single focused cells in each sample. (E-H) mRNA expression of cytokines and transcription factors of *Atf6* WT or KO TH2 (E-F) and TH17 (G-H) cells. (A, B, E-H) *Actb* was used as a loading control for RT-qPCR. (I-J) Western blot of GATA3 and RORγt expression in *Atf6* WT or KO TH2 (I) and TH17 (J) cells, respectively. Protein abundances were normalized to Tubulin. Data are a representative (C-D, I-J left) or a combination (A-B, E-H, C-D right, I-J right) of 3 experiments. Values are means and SD. Student’s *t*-test, * *p* < 0.05. (K) Schematic of a feedforward loop of cytokines-ATF6-mediated UPR-cytokines in TH2 and Th17 cells. Increased ER capacity and cytokine signals also result in increased expression of GATA3 and RORγt in TH2 and TH17 cells, respectively, which further increase cytokine production (not shown).

### ATF6 deficiency alleviates mixed granulocytic airway inflammation

Since ATF6 is induced in asthmatic lungs and highly expressed in TH2 and TH17 cells (Fig. 1A and 1C), we then tested the role of ATF6 in mixed granulocytic airway inflammation, driven by mixed TH2 and TH17 responses. ATF6 KO and control mice were intranasally (i.n.) immunized with HDM-AF (AF extract alone is not sufficient to induce TH2 but mostly TH17 responses in mice on the C57BL/6 background on our hands) and a model antigen chicken ovalbumin (OVA). The frequencies of IL-4^+^ TH2 and IL-17^+^ TH17 were significantly lower in bronchoalveolar lavage fluids (BALFs) of *Atf6* KO mice compared with those of WT mice (Fig. 5A).

**Figure 5.**
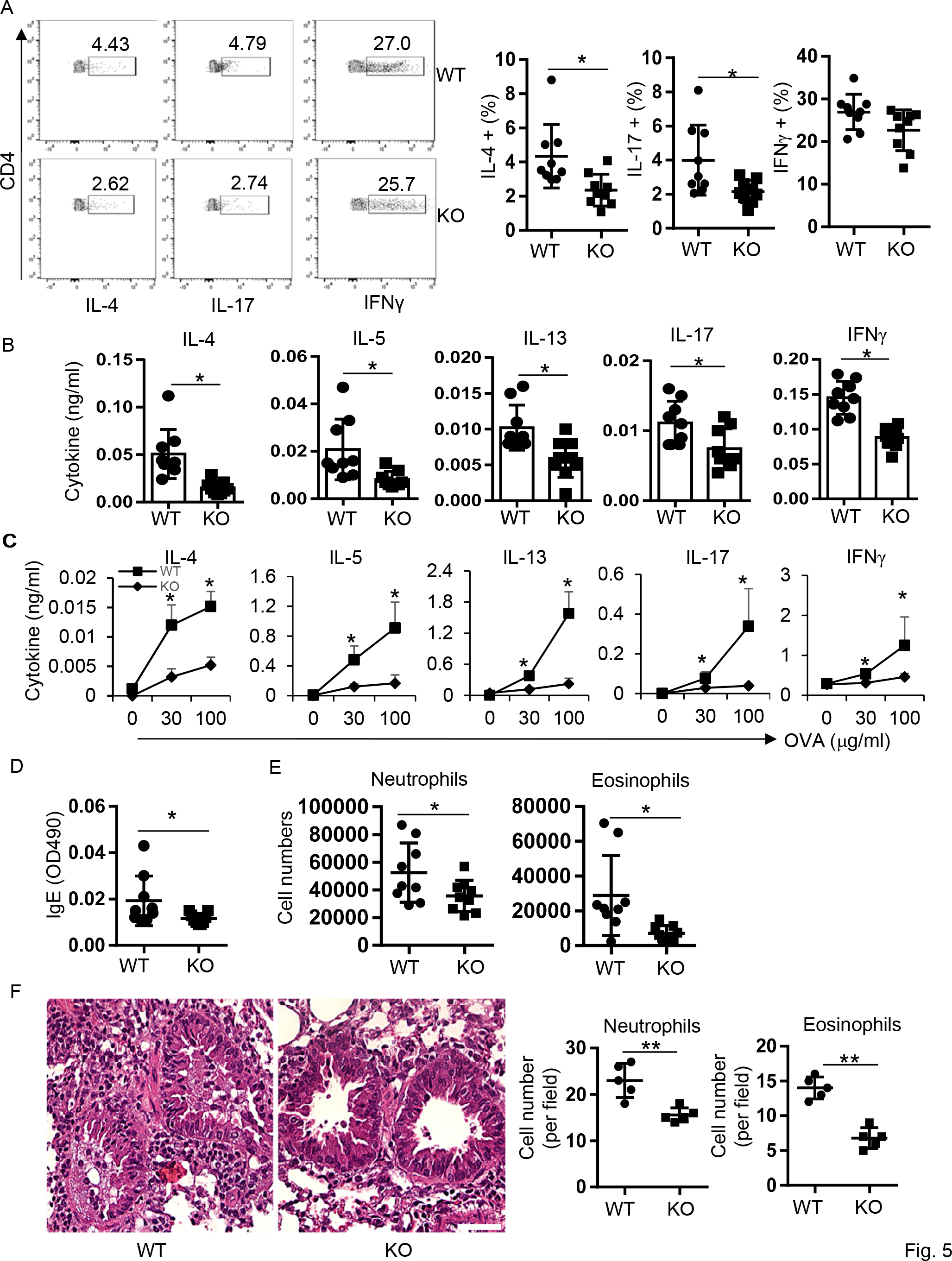
*Atf6*-deficiency alleviates mixed granulocytic airway inflammation. *Atf6* KO and WT mice were subjected to induction of asthma by using HDM-AF-OVA sensitization. (A) Flow cytometry of TH1, TH2, and TH17 cells in BALFs from the asthmatic mice following intracellular staining. (B) ELISA of cytokines in BALFs. (C) ELISA of cytokine expression by asthmatic LLN cells after *ex vivo* recall with OVA at indicated concentrations. (D) ELISA of IgE levels in BALFs. (E) Neutrophils and eosinophils in BALFs. Neutrophils were defined as CD45.2^+^CD11c^−^Ly-6G^+^ whereas eosinophils as CD45.2^+^CD11c^−^Ly- 6G^−^CD11b^+^SiglecF^hi^ by using flow cytometry. (F) Hematoxylin and eosin (H&E) staining of lung sections. Scale bar, 50 μm. Right, statistical analysis of neutrophils and eosinophils in the lung sections. Data (mean ± SD) shown are a representative (A left, C, F) or a combination (A right, B, D-E) of 2 experiments (n = 4-5). Student’s *t*-test, * *p* < 0.05; ** *p* < 0.005.

Consistently, *Atf6* KO BALFs expressed decreased levels of TH2 cytokines IL-4, IL-5, and IL-13, and TH17 cytokine IL-17 relative to WT BALFs (Fig. 5B). Interestingly, *Atf6*-deficiency did not alter the percentage of IFNγ^+^ TH1 cells (Fig. 5A), but impaired the expression of TH1 cytokine IFNγ in the BALFs (Fig. 5B), suggesting that ATF6 influences the cytokine secretion but not the lineage commitment of TH1 cells. Upon *ex vivo* restimulation with various concentrations of OVA, lung-draining mediastinal lymph node (LLN) cells from *Atf6* KO mice expressed lower amounts of TH2 (IL-4, IL-5, and IL-13), TH17 (IL-17), and TH1 (IFNγ) cytokines in comparison with those from WT mice (Fig. 5C). IgE responses are a hallmark of allergic disorders. *Atf6* KO asthmatic mice exhibited decreased levels of OVA-specific IgE than WT mice (Fig. 5D). We next profiled granulocytic airway influxes and found that *Atf6* KO led to significantly fewer neutrophils and eosinophils (Fig. 5E). Histological analyses further revealed that the lungs of *Atf6* KO mice had profound decreases in immune infiltrates, including both neutrophils and eosinophils, at the peribronchial and perivascular spaces than those of WT mice (Fig. 5F). Taken together, ATF6 plays a crucial role in TH cell, especially TH2 and TH17, responses and eosinophil and neutrophil recruitment in allergen-induced airway inflammation.

### Inhibition of ATF6 suppresses both mouse and human memory TH responses

Asthma frequently manifests a chronic phenotype with remission, relapse, and even exacerbation, in which encountering experienced antigens/allergens leads to reactivation of memory TH cells (and mast cells). For instance, in the atopic asthma murine model, memory TH2 cells play an important role in the relapse of disease ^40^. Targeting established TH cells may serve as a therapy for asthma. To test this, we treated murine splenic memory TH cells isolated from WT C57BL/6 mice overnight on an anti-CD3 and anti-CD28 coated plate with a vehicle or various concentrations of ATF6-specific inhibitor Ceapin A7 ^41^. Both treatments with 6 μM and 12 μM Ceapin A7 decreased mRNA expression of ATF6 downstream genes, including *Atf6* itself, compared with the vehicle treatment (Fig. 6A). Administration of Ceapin A7 also significantly downregulated TH2 cytokine IL-5 and TH17 cytokine IL-17 in a dose-dependent manner (Fig. 6B). Similarly, Ceapin A7-treated human peripheral blood CD4^+^CD45RO^+^ T cells expressed decreased mRNA amounts of ATF6 downstream genes and cytokines IL-5 and IL-17 (Fig. 6C- D). These results suggest that ATF6 inhibitor Ceapin A7 suppresses ATF6-mediated UPR and in turn, downregulates both TH2 and TH17 cytokine expression.

**Figure 6.**
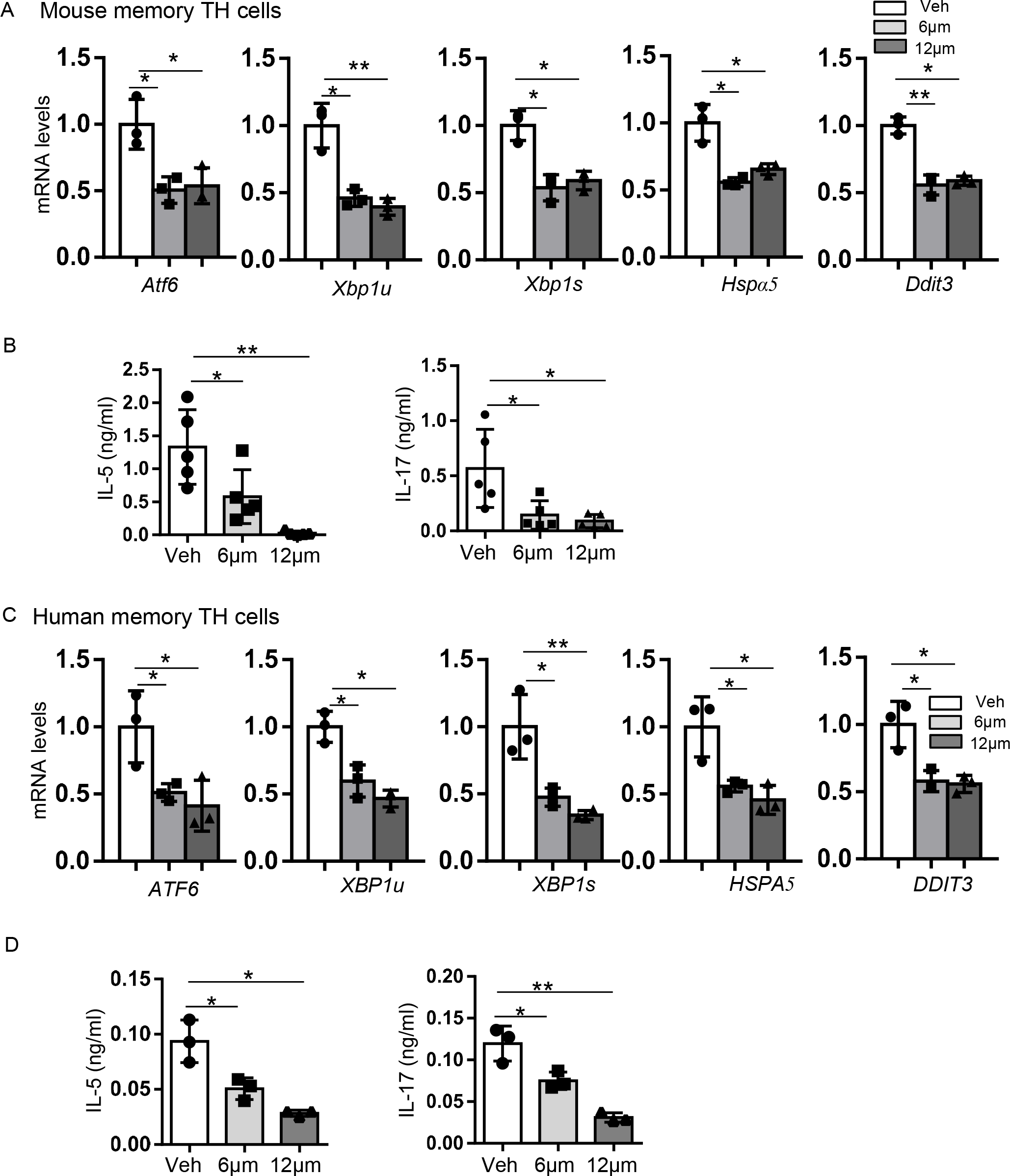
ATF6 inhibitor Ceapin A7 suppresses UPR and cytokine production in mouse and human memory TH cells. (A) RT-qPCR of mRNA expression of ATF6 downstream genes in mouse memory TH cells after treatment with Ceapin A7 at indicated concentrations or a vehicle (Veh). Levels are relative to *Actb*. (B) ELISA of cytokine expression in supernatants of (A). (C) mRNA expression of ATF6 downstream genes in human memory TH cells treated with Ceapin A7 at indicated concentrations or a vehicle. Levels are relative to *ACTB*. (D) ELISA of cytokine expression in supernatants of (C). Data are a combination of 3 (A, C, D) or 5 (B) experiments. Values are means and SD. Student’s *t*-test, * *p* < 0.05, ** *p* < 0.005.

### Inhibition of ATF6 attenuates mixed granulocytic airway inflammation

ATF6 plays an essential role in TH2 and TH17 cell-mediated mix granulocytic asthma (Fig. 5). Since ATF6 inhibitor Ceapin A7 inhibits both murine and human memory TH2 and TH17 responses via suppression of UPR, we then explored whether inhibition of ATF6 *in vivo* benefits mixed granulocytic asthma. As described above, mixed granulocytic asthma was induced in *Atf6* KO and control mice with i.n. administration of HDM-AF with OVA. To avoid its systemic effect, Ceapin A7 (0.5 mg/kg) or a vehicle was intratracheally (i.t.) injected daily since the last two HDM-AF-OVA challenges for 4 days. The mice were analyzed one day after the last dose of Ceapin A7. Administration of Ceapin A7 significantly lessened the frequencies of IL-4^+^ TH2 and IL-17^+^ TH17 but not IFNγ^+^ TH1 cells in BALFs compared with those of vehicle treatment (Fig. 7A). Consistently, BALFs from Ceapin A7-treated mice contained decreased TH2 cytokines IL4, IL-5, and IL-13, and TH17 cytokine IL-17 relative to those from vehicle-treated mice (Fig. 7B). Ceapin A7 treatment did not alter IFNγ levels in BALFs (Fig. 7B). In addition to TH2 and TH17 cytokines, mice receiving Ceapin A7 had fewer neutrophils and eosinophils in the BALFs than those receiving a vehicle (Fig. 7C). In the lung, Ceapin A7 regimen resulted in decreased mRNA expression of neutrophil markers Ly6G and Elane (encoded by *Ela2*) and eosinophil peroxidase (*Epx*) (Fig. 7D). Congruously, lungs from Ceapin A7-treated mice displayed decreased infiltration of immune cells, including neutrophils and eosinophils in the peribronchovascular spaces compared with those from vehicle-treated animals (Fig. 7E). Taken together, ATF6 inhibitor Ceapin A7 suppresses mixed granulocytic airway inflammation, suggesting a potential therapeutic for the mixed type of asthma.

**Figure 7.**
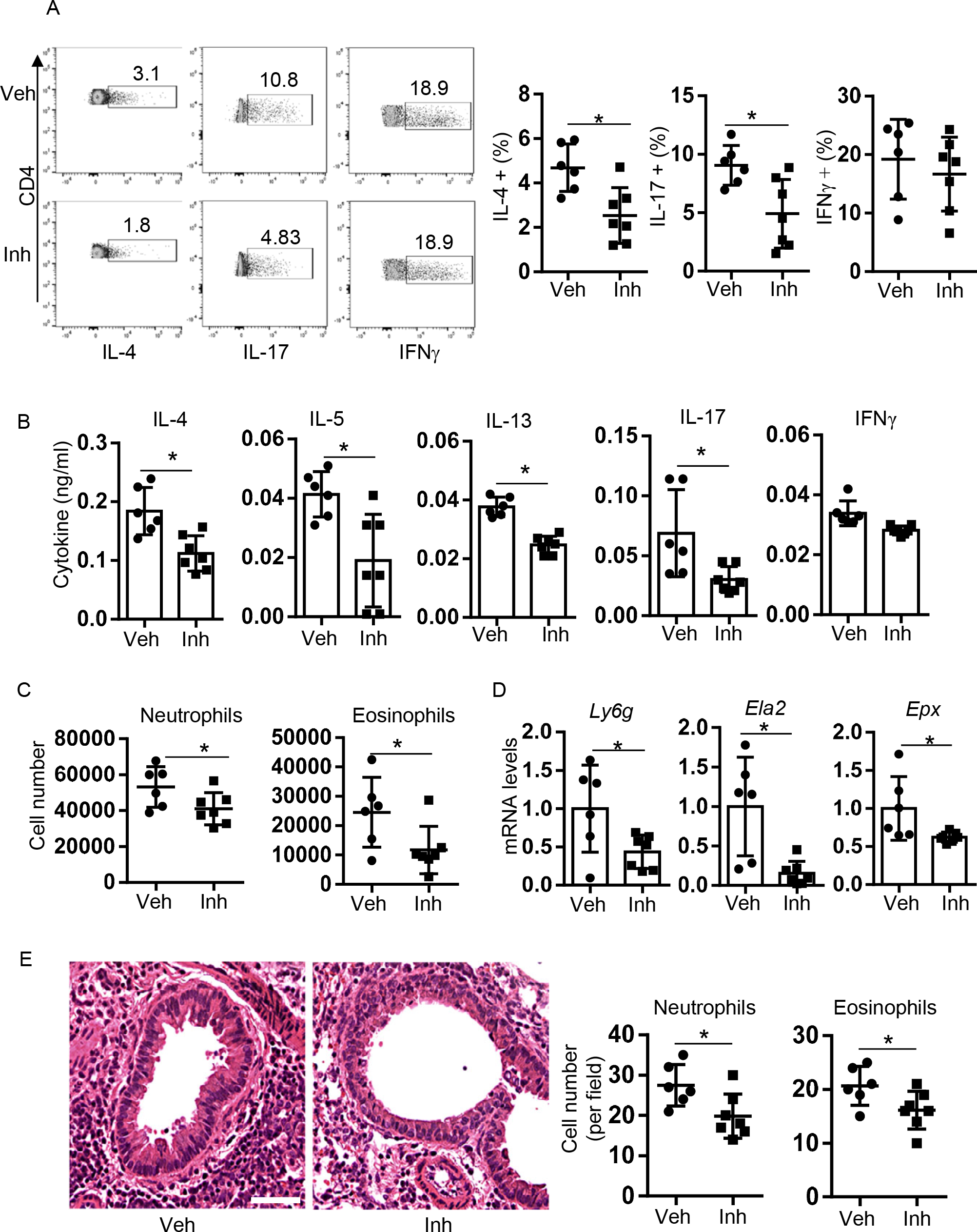
Administration of Ceapin A7 attenuates mixed granulocytic airway inflammation. Mixed granulocytic asthma was induced as described in Fig. 5 and the mice were treated with Ceapin A7 (Inh) or a vehicle (Veh). (A) Flow cytometry of TH1, TH2, and TH17 cells in BALFs following intracellular staining. (B) ELISA of cytokines in BLAFs. (C) Neutrophils and eosinophils in BALFs. (D) RT-qPCR of mRNA expression of *Ly6g* and *Ela2* and *Epx* in asthmatic lungs. (E) H&E staining of lung sections. Scale bar, 50 μm. Right, statistical analysis of neutrophils and eosinophils in the lung sections. Data (mean ± SD) shown a representative of 3 experiments (n = 6-7). Student’s *t*-test, * *p* < 0.05.

## Discussion

UPR is broadly involved in asthmatic responses (see detailed discussion in “Introduction”). We also observed upregulation of UPR in asthmatic mice compared with healthy control mice. As one of the key transducers of UPR, ATF6 plays both protective and deleterious roles in pathogenesis ^20^. Activated ATF6 (ATF6f), acting as a transcription factor, induces a set of genes encoding ER chaperones and enzymes that govern protein folding, secretion, and ER-associated protein degradation, thus resolving ER stress and maintaining intracellular homeostasis ^13–15, 42^. Besides its safeguarding role, ATF6 promotes disease progress via the induction of inflammatory mediators, such as cytokines and chemokines, in immune cells. For instance, genetic ablation of ATF6 leads to decreased expression of IL-17 and IFNγ and amelioration of disease severity in experimental autoimmune encephalomyelitis, a model of human multiple sclerosis ^43^. The pathogenic role of the ORMDL3-ATF6 axis has been characterized in the asthmatic epithelial cells and smooth muscle in humans and mice (see “Introduction”). However, the effects of ATF6 in TH cells are understudied. Our current study showed that ATF6 is highly expressed in TH2 and TH17 cells over TH1 and regulatory T cells, implicating a role of ATF6 in TH2 and/or TH17-driven granulocytic airway inflammation.

Cytokine-mediated activation of STAT proteins plays a pivotal role in the differentiation and maintenance of TH cells ^36, 37^. During naïve CD4^+^ T cell activation, IL-4 signaling activates STAT6 that induces GATA3 and establishes a TH2 lineage program. TH2 cells themselves produce IL-4 that further enforces and maintains the TH2 program via the activation of STAT6. We found that in TH2 cells, the IL-4-STAT6 cascade induces the expression of ATF6. Similarly, IL-6, IL-21, and IL-23, in the context of TGFβ, activate STAT3 that drives TH17 cell polarization. TH17 cell maintenance is mediated by autocrine/paracrine of IL-21 and extracellular IL-23. In TH17 cells, activated STAT3 binds to the *Atf6* gene and induces its expression. Upregulated ATF6 induces UPR-related ER-resident proteins and promotes the secretion of TH signature cytokines. Although ATF6-mediated UPR is a post-transcriptional regulation, it controls the production of functional proteins, including the master transcription factors GATA3 and RORγt in TH2 and TH17 cells, respectively. *In vitro*, *Atf6*-deficiency impairs the differentiation of both TH2 and TH17 cells and fails to induce UPR factors, leading to decreased cytokine production by these cells. Therefore, the cytokine-STAT signaling in TH2 and TH17 cells induces ATF6 that promotes UPR to increase the expression of master transcription factors; these transcription factors further drive the expression of IL-4 and IL-21 in TH2 and TH17 cells, respectively, which through activation of STAT6 and STAT3, respectively, form a feedforward loop (Fig. 4K), heightening TH2 and/or TH17 effector function and potentially increasing their pathogenesis.

To gain an insight into the T cell-intrinsic role of ATF6 in airway inflammation, we employed an HDM-AF caused mixed granulocytic asthma model. Mice bearing *Atf6*-deficient T cells had much milder airway eosinophilia and neutrophilia with fewer immune infiltrates in the lungs and decreased frequencies of TH2 and TH17 cells in the BALF. Collectively, these results demonstrate that ATF6 is a crucial factor regulating not only airway structural cell hyperplasia ^32, 33^, but also TH2 and TH17 responses and associated eosinophil and neutrophil recruitment in allergen-elicited airway inflammation.

Patients with severe asthma respond poorly to current therapies, including corticosteroids even combined with bronchodilators ^4, 5, 44^. In a mouse model, adoptive transfer of TH17 but not TH2 cells leads to a corticosteroid dexamethasone-resistant airway phenotype with increased airway hyperresponsiveness in an IL-17-IL17R signal-dependent manner ^45^, highlighting the importance of the TH17-IL-17 axis in treatment-refractory severe asthma. In another study, blockade of IL-4 and/or IL-13 enhances TH17 responses and airway neutrophilia, whereas neutralization of IL-13 and IL-17 diminished both airway eosinophilia and neutrophilia and associated mucus hyperplasia and airway hyperreactivity, suggesting that simultaneously targeting both pathways may maximize therapeutic efficacy in patients manifesting both TH2 and TH17 endotypes ^46^.

Since ATF6 promotes both TH2 and TH17 responses and enhances mixed granulocytic airway disease, targeting ATF6 may benefit patients with mixed granulocytic asthma or TH2 or TH17 biased asthma. Ceapin A7 is a recently developed ATF6-specific inhibitor ^41^. We found that *in vitro*, administration of Ceapin A7 suppresses the expression of ATF6 downstream genes that mediate UPR and diminishes the expression of TH2 and TH17 cytokines by murine memory TH cells. Moreover, in a mixed granulocytic asthma model, we observed that treatment with Ceapin A7 at the chronic state represses both TH2 and TH17 responses, resulting in decreased airway eosinophilia and neutrophilia and attenuated lung inflammation including decreased eosinophil and neutrophil infiltration in the peribronchovascular interstitium. Our results demonstrate primitive efficacy of ATF6 inhibitor Ceapin A7 in the treatment of mixed granulocytic asthma. In addition, treatment with Ceapin A7 inhibits ATF6-mediated UPR and decreases the expression of TH2 and TH17 cytokines in human memory TH cells, similar to those observed in murine memory TH cells, implicating a promising treatment by blockade of ATF6. Whether this inhibitor can alleviate human asthma, especially T2-low or mixed granulocytic severe asthma, is to be tested.

In conclusion, our results demonstrate that cytokine signals induce ATF6 in TH2 and TH17 cells and ATF6 boosts TH2 and TH17 responses via UPR. These together form a feedforward circuit in TH2 and TH17 cells, promoting mixed granulocytic airway disease. Targeting ATF6 may serve as a potential therapy for mixed and even T2-low endotypes of severe asthma.

## Materials and Methods

### Induction of mixed granulocytic asthma

For induction of allergic asthma, 5–8-week old *Atf6*^fl/fl;CD4Cre^ (*Atf6* KO) and *Atf6*^fl/fl^ (*Atf6* WT) littermates were immunized intranasally (i.n.) with 100 μg HDM and 50 μg *A. fumigatus* (AF)-extracts with 50 μg OVA in 30 μl PBS on days 0, 2, 7, and 14 and day 15. On day 16, BALFs were collected for analysis of airway infiltrates, and LLNs and lungs were immediately collected and stored on ice for analysis of immune responses and immune infiltration, respectively. The superior lung lobe was fixed in 2% paraformaldehyde for histology and the other lobes were meshed in 1 ml PBS with 120-micron nylon mesh. The lung suspension was spun to obtain supernatant for ELISA measurement of cytokines, and the cell pellet was treated with red cell lysis butter (0.1 mM EDTA, 150 mM NH4Cl, and10 mM KHCO3) to remove red cells. A small portion (5%) of the lung single-cell suspension was used for RNA preparation. All mice were housed in the specific pathogen-free animal facility and animal experiments were performed with protocols approved by the Institutional Animal Care and Use Committee of the University of New Mexico Health Sciences Center.

### TH1, TH2, and TH17 cell differentiation

CD4^+^CD25^−^CD44^−^CD62L^+^ naïve T cells were isolated from ATF6 WT and KO mice and differentiated in a TH1 (10 ng ml^−1^ IL-12 and 5 μg ml^−1^ anti-IL4), TH2 (5 ng ml^−1^ IL-4 and 5 μg ml^−1^ anti-IFNγ), or TH17 (2 ng ml^−1^ TGFβ, 10 ng ml^−1^ IL-6, 2 μg ml^−1^ anti-IFNγ, and 2 μg ml^−1^ anti-IL4)-polarizing condition on an anti-CD3 and anti-CD28 (1 μg ml^−1^, each) coated plate.

### Overexpression of constitutively active STAT6 and STAT3

*In vitro* differentiated TH2 and TH17 cells were rested for 24 h on a plate without coating anti-CD3 and anti-CD28 on day 4. On day 5, the rested TH2 and TH17 cells were transfected with STAT6VT in pMX-IRES-hCD2 vector ^47, 48^ and STAT3C in pGFP-RV vector ^49, 50^, respectively, with a corresponding control vector by electroporation with a Neon transfection system (1600v, 20 mS/pulse, 2 pulses; Thermo Fisher Scientific) and incubated on an anti-CD3/anti-CD28 coated plate for 48 h.

Afterward, human CD2^+^ or GFP^+^ cells were sorted from the resulting cultures and subjected to RT-qPCR.

### Treatment with ATF6 inhibitor *in vitro*

Mouse memory CD4^+^ T cells were enriched from WT C57BL/6 splenocytes by negatively depletion of B220^+^ B cells, CD8^+^ T cells and CD25^+^ regulatory T cells and then CD62L^+^ naïve T cells using magnetic beads. Human peripheral blood CD4^+^CD45RO^+^ T cells were purchased from STEMCELL (#70031). Human and mouse memory CD4^+^ T cells were treated overnight with ATF6 inhibitor Ceapin A7 (2323027-38-7, Milliporesigma) at indicated concentrations or a vehicle (10% DMSO-PBS). The supernatants were used for cytokine expression assays and cells for mRNA expression assays.

### Treatment with ATF6 inhibitor *in vivo*

Asthma was induced using HDM-AF with OVA as described above. On days 15-17, the mice i.t. received daily a dose of Ceapin A7 (0.5 mg kg^−1^) or vehicle (10% DMSO-PBS). On day 18, the resulting mice were analyzed as described above.

### Flow cytometry

Antibodies against CD4 (RM4.5), CD11b (M1/70), CD11c (N418), Ly-6G (1AB-Ly6g), SiglecF (E50-2440), Gr-1 (RB6-8C5), CD45.2 (104), IL-13 (eBio13A), IFNγ (XMG1.2), IL-17 (eBio17B7), Foxp3 (FJK-16s) were purchased from eBioscience. IL-4 (11B11) and IL-5 (TRFK5) were from Biolegend. Following surface and/or intracellular stain, cells were analyzed on an Attune NxT Flow Cytometer (Thermo Fisher Scientific) and data were processed by Flojo software v10.5.0 or later (FlowJo, LLC).

### Imaging flow cytometry

In vitro-differentiated TH2 and TH17 cells were fixed with 4% paraformaldehyde at 4℃ for 15 min. The cells were permeabilized and stained with antibodies against CD4, IL-4 or IL-17, Calr (A-9, Santa Cruz Biotechnology), and DAPI (D1306, Fisher Scientific). Single cell images were acquired with Amnis® ImageStream® X MkII (Luminex). Collected data were analyzed with IDEAS 5.0 software. Single cells were determined by the area and aspect ratio and focused cells by the gradient root mean square (RMS) feature. The relative brightness of a given fluorochrome on a cell was quantified by sum of the pixel intensities within a given channel mask normalized to the camera dark-current. A total of 1000 CD4^+^ IL-4^+^ (or IL- 17^+^) single focused cells each sample were analyzed.

### Western blot

Cell lysates were prepared by using Laemmli’s sample buffer. After being reduced with 2.5% β-mercaptoethanol at 100 °C for 5 min, the lysates were resolved on SDS-PAGE gels and blotted with indicated antibodies following a standard protocol. Immunoblot antibodies were anti-ATF6 (A12570, ABclonal), anti-Xbp1s (ab220783, Abcam), anti-GATA3 (16E10A23, Biolegend), anti-RORγt (B2D, eBioscience), and anti-α-Tubulin (eBioP4D1, eBioscience).

### ELISA

LLN cells (4 × 10^6^ cells ml^−1^) from asthmatic mice were recalled with various concentrations of OVA for 3 days and the supernatants were collected for measurement of cytokine expression by ELISA using a standard protocol. For *in vitro* differentiated TH1, TH2, and TH17 cells, the supernatants were collected after restimulation overnight on an anti-CD3 plate and used to measure cytokine expression by ELISA. In some experiments, BALF or lung homogenate supernatants were to measure cytokines. To measure OVA-specific IgE, plate-bound OVA (100 μg ml^−1^) was used as capture following detection with anti-IgE (23G3, eBioscience) as a standard protocol.

### RT-quantitative (q) PCR

Gene mRNA expression was determined by qPCR with a CFX Connect Real-Time PCR Detection System (Bio-Rad Laboratories) following reverse transcription with SuperScript Reverse Transcriptase (18064022, Thermo Fisher Scientific). Data were normalized to a reference gene *Actb* (or *ACTB*). The primers of mouse genes were: *Atf6* forward, 5’-AGAGTCTGCTTGTCAGTCGC, reverse, 5’-GCAGGGCTCACACTAGGTTT; *Il21* forward, 5’-ACAAGATGTAAAGGGGCACTGTGA, reverse, 5’-ACAGGAGGTACCCGGACACA; and *Actb*, *Gata3*, *Il4*, *Il5*, *Il13*, *Xbp1s*, *Hspa5*, and *Ddit3* ^51^, *Ela2* and *Epx* (*Epo*) ^52^, *Il17* ^53^, *Il17f* ^54^, *Xbp1u* ^55^, *Rorc(gt)* ^56^, and *Ly6g* ^57^ as described. The primers of human genes were: *ACTB* forward, 5’-GACGGCCAGGTCATCACCATTG, reverse, 5’-GTACTTGCGCTCAGGAGGAG; *ATF6* forward, 5’-CCACTAGTAGTATCAGCAGGAAC, reverse, 5′-GGTAGCTGGTAACAGCAGGT; *HSPA5* forward, 5’- AAGCTGTAGCGTATGGTGCT, reverse, 5’-TCAGGGGTCTTTCACCTTCAT; and *DDIT3* forward, 5’-TGGAACCTGAGGAGAGAGTGTT, reverse, 5’- GGATAATGGGGAGTGGCTGG; *and XBP1u* and *XBP1s* as described ^58^.

### Chromatin immunoprecipitation (ChIP)

Cell lysates were prepared as described ^59^. The lysates were pre-cleared with Protein G magnetic beads at 4 °C for 30 m. After removal of the Protein G beads, supernatants were collected and incubated with an antibody against the target protein or a control antibody overnight at 4 °C, followed by incubation with Protein G magnetic beads for 2 h at 4 °C the next morning. The bead-antibody-chromatin complexes were magnetically separated and washed twice with low salt TE buffer and additional twice with high salt TE buffer. Finally, the bead-antibody-chromatin complexes were collected and resuspended in TE buffer for reverse crosslink in the presence of 0.2 M NaCl for 4 h at 65 °C. The samples were then subjected to qPCR. The antibodies were pSTAT6 (D8S9Y, Cell Signaling) and pSTAT3 (B-7, Santa Cruz). STAT6 ChIP primers were: Site 1 forward, 5’- TTCCGCAATCGCAATCGGTA, reverse, 5’-GACCGAGAGGAACTTTTCCCA; Site 2+3 forward, 5’-CGGAGGATTCTGGGAAGCAT, reverse, 5’-AGACGGTGAATGAACACGGA; Sites 4 forward, 5’-GTAAGTGCGGACATGGAGAA, reverse, 5’- GACATGTGTTAGAACCACTCCT; Site 5 forward, 5’-ACCTCGGATTCTGTCCCAGT, reverse, 5’-ACGATATAGACATCTCAGAACGAAA; Sites 6+7 forward, 5’- AAAACCCCACAGCGGGAC, reverse, 5’-CGGAGGCAAACTATCTCACCG; Site 8 forward, 5’-TGGATCTGCGAATGTACCCG, reverse, 5’-CTTCTACGACCCTCAACCCG, and Sites 9+10 forward, 5’-GTCCTGGCCGTAGCAATAAT, reverse, 5’-GGCAGCTACAATCATCATCCT. STAT3 ChIP primers were: Site 7 forward, 5’- ACGGGTTGAGGGTCGTAGAA, reverse, 5’-CCAGGAACACACGCTGAAAAA, and Sites 1+2, 3+4, 5, 6, and 8 primers as same as STAT6 Sites 1, 2+3, 6+7, 8, and 9+10, respectively.

### Statistical analysis

Results were expressed as mean ± SD. Differences between groups were calculated for statistical significance using the unpaired Student’s *t*-test. *P* ≤ 0.05 was considered significant.

### Data Availability

All data generated or analyzed during this study are included in this article.

## Acknowledgments

This work was supported in part by NIH grants HL148337 and AI142200 (X.O.Y.), and DK110439 and DK132643 (M.L.). We acknowledge the University of New Mexico Comprehensive Cancer Center Flow Cytometry Core supported by NIH CA118100 and the Autophagy, Inflammation, and Metabolism in Disease Center Inflammation and Metabolism Core supported by NIH GM121176.

## Author Contributions

X.O.Y., M.L., and X.W. designed the study and coordinated the experiments; D.W., X.Z., K.Z., R.W., C.W., and X.O.Y. performed the experiments; D.W. and X.O.Y. wrote and all authors reviewed the manuscript.

